# Lack of motor defects and ALS-like neuropathology in heterozygous *Sptlc1* Exon 2 deletion mice

**DOI:** 10.1101/2025.02.18.638951

**Authors:** Devesh C. Pant, Museer A. Lone, Janani Parameswaran, Fuying Ma, Prisha Dutta, Zitong Wang, Jaekeun Park, Sumit Verma, Thorsten Hornemann, Jie Jiang

## Abstract

Mutations in the human *SPTLC1* gene have recently been linked to early onset amyotrophic lateral sclerosis (ALS), characterized by global atrophy, motor impairments, and symptoms such as tongue fasciculations. All known ALS*-*linked *SPTLC1* mutations cluster within exon 2 and a specific variant, c.58G>T, results in exon 2 skipping. However, it is unclear how the exon 2 deletion affects SPTLC1 function *in vivo* and contributes to ALS pathogenesis. Leveraging the high genomic sequence similarity between mouse and human *SPTLC1*, we created a novel mouse model with a CRISPR/Cas9-mediated deletion of exon 2 in the endogenous murine *Sptlc1* locus. While heterozygous mice did not develop motor defects or ALS-like neuropathology, homozygous mutants died prematurely. These findings indicate that *Sptlc1* ΔExon2 heterozygous mice do not replicate the disease phenotype but provide valuable insights into SPTLC1 biology and serve as a useful resource for future mechanistic studies.

## Introduction

Early-onset ALS, also known as juvenile ALS (jALS), is a rare subtype of ALS that affects individuals under the age of 25. While ALS primarily manifests in individuals aged 50-70, studying jALS can provide valuable insights into ALS pathogenesis, as both forms share the hallmark degeneration of upper and lower motor neurons. The clinical heterogeneity of jALS poses significant challenges for diagnosis and management, with many patients remaining undiagnosed and the underlying causes of their symptoms elusive. Advances in sequencing technologies have uncovered novel genetic defects in ALS that were previously unrecognized as contributors to neurological diseases [1]. Recently, *SPTLC1* (OMIM# 605712) variants have been reported in jALS cohorts across European and Asian populations [2–6], joining a growing list of genes associated with jALS, including FUS, SETX, ALS2. Electrophysiological and histopathological analyses of muscle and nerve biopsies from SPTLC1-jALS patients reveal diffuse acute and chronic denervation in multiple myotomes without sensory neuropathy. Notably, mild cognitive dysfunction has been reported in two SPTLC1-jALS patients, reinforcing the idea that ALS forms a disease spectrum with dementia [3].

SPTLC1 is a highly conserved gene that encodes a subunit of serine palmitoyltransferase (SPT), the enzyme catalyzing the first step in sphingolipid biosynthesis. Sphingolipids are critical components of cell membranes and signaling pathways, and their involvement in central nervous system (CNS) disorders has been discussed in our review [7]. The SPT complex, a multi-subunit enzyme, conjugates palmitoyl-CoA and L-serine to produce long-chain bases, the precursors of sphingolipids. The SPT enzyme complex resides in the endoplasmic reticulum (ER) membrane and at the ER-mitochondria contact sites. In mammals, the enzyme consists of two of three primary subunits—SPTLC1/SPTLC2 or SPTLC1/SPTLC3 —that assemble into tetrameric complexes [8, 9]. Mutations in the C-terminus of *SPTLC1* have been previously linked to hereditary sensory and autonomic neuropathy type 1 (HSAN1), a group of rare peripheral neuropathies characterized by the formation of non-canonical deoxy-sphingolipids [10, 11]. In contrast, ALS-associated *SPTLC1* variants are primarily found near the transmembrane (TM) interface encoded by exon 2. One variant (*SPTLC1* c.58 G >T) results in a single base-pair change adjacent to the splice acceptor site for exon 2. Instead of the predicted amino acid substitution, the c.58G>T variant predominantly leads to an in-frame deletion of amino acids 20-56 comprising the entire TM domain.

The *SPTLC1* gene is ubiquitous which initiates *de novo* sphingolipid synthesis. SPTLC1 linked HSAN1 mutation (C133W) have been previously modeled in mice using transgenic or knock-in approaches [12, 13]. The transgenic mice were reported to display mild phenotype. However, the knock-in heterozygotes were reported normal and homozygous die embryonically [14]. To date, there are no published murine models for SPTLC1-related ALS. The current evidence surrounding sphingolipid profiles in SPTLC1 ΔExon2 ALS human patients remains sparse. One study indicated an increase in serum sphingolipids [5], whereas another group reported no changes in plasma sphingolipids (ceramides, sphingomyelins) levels of exon2 skipping variant in human patients [3]. It is to be noted that blood lipids can vary considerably within individuals over short periods of time due to intrinsic factors, such as hormonal variation or diet [15]. Given the very small sample sizes in original studies, it’s difficult to draw meaningful insights from these as well as from follow-up overexpression studies using cell lines [3, 5, 16].

In this study, we generated *Sptlc1* exon 2 deletion (ΔE2) knock-in mice to investigate the pathogenic mechanisms of *SPTLC1*-linked ALS. Mice lacking *Sptlc1* exon 2 in both alleles exhibited embryonic lethality with incomplete penetrance. Longitudinal motor function assessment revealed that the SPTLC1 ΔE2 mutation did not induce an ALS-like phenotype in heterozygous mice, nor did it alter canonical sphingolipid levels. Furthermore, neuropathological analysis showed no changes in markers of astrogliosis or microglial activation. Our findings provide new insight into the *in vivo* roles of SPTLC1 ΔE2 mutation and suggest that the ΔE2 heterozygous mutation is insufficient to induce an ALS-like phenotype in mice.

## Materials and methods

### Generation of the *Sptlc1^ΔE2^* mouse model

The *Sptlc1* knock-in mouse model, bearing an exon 2 deletion (ΔE2) in the *Sptlc1* gene (ENSMUSG00000021468), was generated by CRISPR/Cas9 genome editing. Both mouse and human SPTLC1 are highly conserved (**Fig. S1**). Zygotes from C57BL/6J (RRID: IMSR_JAX:000664) mice were microinjected with guides (sgRNAs) targeting the introns flanking exon 2 (GCAGACTTTGAAGCAGCTTC, TTGTTTCACTCGAGGTATGT, GAAAGGCTTGAGAGCCATTG), Cas9 protein (Synthego, CA) and 120 bp donor (GTGTGGTAAGAATGTGAAGGTCAGGAGAGGCAGCACTGTGGTGGGCTCCATTTCCCAGCATGTTGGTGCTACATCTCAGAAGGACTCAAACTGGACCCTGTAGTGTGTCAAAATGCTTCA). These zygotes were then subsequently implanted into surrogate female mice with the assistance of Emory Mouse Transgenic and Gene Targeting Core. All mouse experimental protocols were approved by the Institutional Animal Care and Use Committee at Emory University. Mice were housed on a 12/12 h light/dark cycle with access to standard mouse chow (LabDiet Cat# 5001) and water *ad libitum*. Two founder mice (F_0_) with a 195 bp deletion encompassing exon 2 were identified and selected for colony establishment (**Fig. S2**). To minimize mosaicism and potential off-target events, F_0_ mice were backcrossed with wild type C57BL/6J mice before generating experimental cohorts described. Genotyping using genomic DNA was performed using the following primers: GCTGACAGTGTTGGGGTTTT, ACTTCGTACCCCCAGCTCTC. The in-frame deletion of exon 2 was confirmed by polymerase chain reaction (PCR) using cDNA from heterozygous mice by using primers in exon 1 and exon 5 and exon 2 deletion was further validation by sanger sequencing (**Fig. S3**).

### Mouse behavioral testing

Experimenters were blinded to the genotype of the mice for all experiments. *Sptlc1^+/+^*, *Sptlc1^ΔE2/+^* mice were used in all behavioral tests. Mice were habituated to the testing room for 30-45 minutes (unless otherwise specified) prior to the start of each behavioral experiment to minimize stress-induced variability.

### Grip strength assessment

Forelimb and hindlimb grip strength were assessed using a Grip Strength Meter (Ugo-Basile). The apparatus consists of a base plate and a T-shaped grasping bar with adjustable height, attached to a force transducer connected to a peak amplifier. Before each measurement, the gauge was reset to 0 g after stabilization to ensure accuracy. The mouse was placed over the base plate in front of the bar and its tail was slowly pulled back by the experimenter until the animal released its grip. The maximal tension recorded at the time of release was used as the grip strength.

### Rotarod test

Locomotor function was assessed using a rotarod apparatus (Ugo-Basile) with a gradually increasing speed protocol. The speed ramped from 5 to 40 rpm over 5 minutes, with a maximum trial duration of 10 minutes. Mice underwent 3 trials per day for 3 consecutive days, with a 60-minute acclimation period in the testing room before the first trial each day. During each trial, the latency to fall from the rotating rod or to make two consecutive complete rotations while clinging to the rod was recorded. Mice were allowed to rest for at least 15 minutes between each trial. The average latency across the three trials was calculated as an indicator of locomotor function. Between runs, the rotarod apparatus was cleaned with 70% ethanol to maintain consistent testing conditions.

### Compound muscle action potential recording

Electrophysiological measurements of compound muscle action potential (CMAP) were performed using needle electrodes, a minimally invasive and highly sensitive method for monitoring neuromuscular function, as previously described [17]. Briefly, mice were anesthetized with isoflurane (3–5% for induction, 2–3% for maintenance) to ensure immobilization and minimize discomfort. Monopolar needle electrodes (Rhythmlink, Columbia, SC) were placed in the left hindlimb for motor nerve stimulation and CMAP recordings. An active needle electrode was inserted in the left gastrocnemius-soleus muscle, while the reference was inserted in the ipsilateral tendinous heel. The ground electrode was inserted subcutaneously in the upper midback of the mouse. The cathode and anode needle electrodes were inserted on either side of the left sciatic nerve at the proximal thigh. The left sciatic nerve was stimulated with a 0.1 ms pulse duration and an intensity 1–10 mA using portable electrodiagnostic system (Cadwell Sierra Summit, Kennewick, WA). CMAP peak-to-peak amplitudes were recorded three times while increasing current intensity and adjusting the active recording electrode to obtain a supramaximal CMAP. The highest amplitude among the three recordings was used for data analysis.

### *In vivo* magnetic resonance imaging

Magnetic resonance imaging (MRI) of mouse brains was performed using a 9.4T/20 cm Bruker Avance NEO system in collaboration with the Emory Center for Systems Imaging Core. A two-coil actively decoupled imaging set-up was employed, consisting of a 3 cm ID surface coil for reception and a quadrature coil for transmission. Mice were anesthetized with 2% isoflurane and positioned using an animal holding system. Breathing rate was continuously monitored throughout the procedure, and body temperature was maintained at 37°C using a heated circulating water system. Temperature was monitored via a rectal fiber optic probe to ensure physiological stability during imaging. Coronal T2-weighted images were acquired with a Rapid Acquisition with Refocused Echoes (RARE) sequence. The imaging parameters were as follows: Repetition time (TR) = 4000 ms, Effective Echo time (Eff.TE) = 42 ms, RARE factor = 8, Feld of view = 20 × 20 mm^2^, Matrix = 116 × 116, Average (Avg) = 12, slice thickness = 0.75 mm, number of slices = 11. MRI acquisition was conducted using the ParaVision software (Bruker), and the resulting 2dseq format files were converted to digital imaging and communications in medicine (DICOM) format for downstream analysis. Whole brain, cortex, and brainstem regions were segmented and analyzed using the 3D Slicer image computing platform [18].

### Lipidomics

Lipids were extracted from frozen tissues with 500 μl of methanol containing 200 pmoles of the following internal standards: D_7_-sphinganine (d18:0), D_7_-sphingosine (d18:1), dihydroceramide (d18:0/12:0), ceramide (d18:1/12:0). Samples were incubated on a shaker-incubator at 37°C and 1400 rpm for 1 hour. Lipids were hydrolyzed overnight at 65°C and subsequently re-extracted with chloroform as described previously [12]. Hydrolyzed lipids were dried under a stream of nitrogen gas and resuspended in 200 μl (tissues) or 70 μl (cells) of reconstitution buffer (70% methanol, 10 mM ammonium acetate, pH 8.5). Long chain bases (LCBs) were separated via a reverse-phase C18 column (Uptispere 120 Å, 5 μm, 125 × 2 mm, Interchim, France) connected to a QTRAP 6500+ LC-MS/MS System (Sciex). LCBs were quantified by normalization to the internal standards and either plasma volume or protein content for tissues. For protein normalization, tissue pellets were homogenized in 8M urea in a Precellys 24 tissue homogenizer (Bertin Technologies) post extraction. Protein concentration was determined using the Bradford assay.

### Immunofluorescence staining

Mouse brain and spinal cord tissues were fixed in 4% paraformaldehyde (PFA) for 24 hours at 4°C and cryoprotected in 30% sucrose (Sigma) for 48 hours. The tissues were embedded in optimal cutting temperature (OCT) compound and sectioned at 20 µM thickness. Sections were mounted on Superfrost Plus glass slides (VWR) and stored at −20 °C. Sections were permeabilized with 0.2% Trition X-100 (Sigma) for 10 minutes and blocked in 5% bovine serum albumin (BSA) for 1 hour at room temperature. Primary antibody diluted in blocking solution were applied and sections were incubated for 24 hours at 4 °C. The following primary antibodies were used: anti-IBA1 antibody (rabbit polyclonal, Wako-01919741, 1:1000) and anti-GFAP antibody (rabbit polyclonal, ab7260, 1:500). Sections were rinsed with 1× PBS and then incubated with secondary antibody (488 Alexa Fluor donkey anti-rabbit IgG (H + L), 1:500 from ThermoFisher Scientific) and DAPI (Millipore D6210, 1:1000) for 24 hours at 4 °C. Sections were rinsed with PBS and mounted with Prolong Gold Antifade mountant (ThermoFisher Scientific, P36930). Imaging was performed using a Keyence BZ-X819 microscope, and images were analyzed by ImageJ software.

### Western blotting

Whole cell extracts were prepared using RIPA Lysis Buffer (pH 7.4, Bio-world, USA) supplemented with Halt^TM^ protease and phosphatase inhibitor cocktail (ThermoFisher Scientific). DNA shearing was performed to reduce viscosity. After centrifugation, proteins concentrations were determined using the BCA Protein Assay Reagent (Pierce, USA). Twenty microgram of protein was resolved on a 4–20% precast polyacrylamide gel (Bio-Rad, USA), and transferred to nitrocellulose membranes (Bio-Rad). Membranes were blocked and incubated overnight at 4 °C with primary antibodies: mouse anti-SPTLC1 (1:2000; BD Biosciences), rabbit anti-β actin (1:2000; GeneTex), and rabbit anti-GAPDH (1:2000; Cell Signaling Technology). Membranes were then incubated for 1 hour at room temperature with secondary antibodies: HRP-conjugated secondary antibodies (ABclonal) or IRDye secondary antibodies (Li-Cor). For detection, Super Signal West Pico (Pierce, USA) was used to visualize peroxidase activity. Protein molecular masses were determined by comparison with protein standards (ThermoFisher Scientific). Band intensities were quantified using ImageJ software and normalized to GAPDH or β-actin as loading controls.

### RT-PCR analysis

Total RNA was prepared from cells using Trizol reagent (Invitrogen) and purified with the quick-RNA miniprep kit (Zymo Research) according to the manufacturer’s instructions. Complementary DNA (cDNA) was generated using high-capacity cDNA reverse transcription Kit (ThermoFisher Scientific) following the manufacturer’s protocol. PCR amplification was performed under the following conditions: 95°C for 30 seconds, (95°C for 30 seconds, 57°C for 30 seconds, 72°C for 1 minute) x 35 cycles, followed by an extension stage of 72°C for 5 minutes and a 10°C hold. To quantify relative mRNA expression, qPCR was performed using Taqman gene expression assays (FAM-Labeled, ThermoFisher Scientific) on a Quantstudio 6 Flex system (Applied Biosystems). The following Taqman assay probes were used: *Sptlc1* (Mm00447343_m1, Exon 3-4), *Sptlc2* (Mm00448871_m1, Exon 4-5), *Ormdl3* (Mm00787910_sH, Exon 4), *Iba1* (Mm00479862_g1, Exon 4-5), *Gfap* (Mm01253033_m1, Exon 6-7). Relative RNA expression was normalized to *Gapdh* (Mm99999915_g1, Exon 2-3).

### Statistical analysis

All the statistical analyses and graphs were prepared in GraphPad Prism (version 9). Data are expressed as mean ± SD or mean ± SEM as indicated in the respective figure legends. Student’s t-test was used for two-group comparisons, and one-way ANOVA was used for multiple group comparisons unless otherwise specified in the figure legends. A *p*-value less than 0.05 was considered statistically significant.

## Results

### Generation of *Sptlc1* exon 2 deletion knock-in mouse model

Exon 2 represents the most common mutational hotspot in human SPTLC1-associated jALS. To develop an *in vivo* model that closely mirrors the human condition and to investigate the underlying disease mechanism, we generated the first *Sptlc1* knock-in mouse model harboring an exon 2 deletion (ΔE2) using CRISPR-Cas9 genome editing (**Fig. 1A**). Two founder lines carrying a 195 bp deletion encompassing exon 2 were established. PCR amplification of genomic DNA followed by agarose gel electrophoresis revealed distinct band patterns: heterozygous *Sptlc1^ΔE2/+^* mice had two bands of 434bp and 239bp respectively, the difference corresponding to the deletion of exon 2; wild type *Sptlc1^+/+^* mice displayed a single 434bp band; and homozygous *Sptlc1^ΔE2/ΔE2^* mice displayed one single band of 239bp (**Fig. 1B**). Sanger sequencing of the PCR products confirmed the removal of exon 2 in these mice (**Fig. 1C**). Among over 100 offspring from 10 independent matings, only 3 homozygous mutant mice were identified, compared to 33 wildtype and 64 heterozygous littermates. These 3 homozygous animals ultimately failed to thrive, dying within 4 weeks of age, suggesting mice with homozygous ΔE2 are embryonic lethal with incomplete penetrance. To assess protein levels, spinal cord tissues were collected from 3-week-old animals. Western blot analysis showed markedly reduced SPTLC1 protein levels in homozygous mutants, while heterozygous *Sptlc1*^ΔE2/+^ mice exhibited no significant changes compared to controls (**Fig. 1D**).

**Figure 1:**
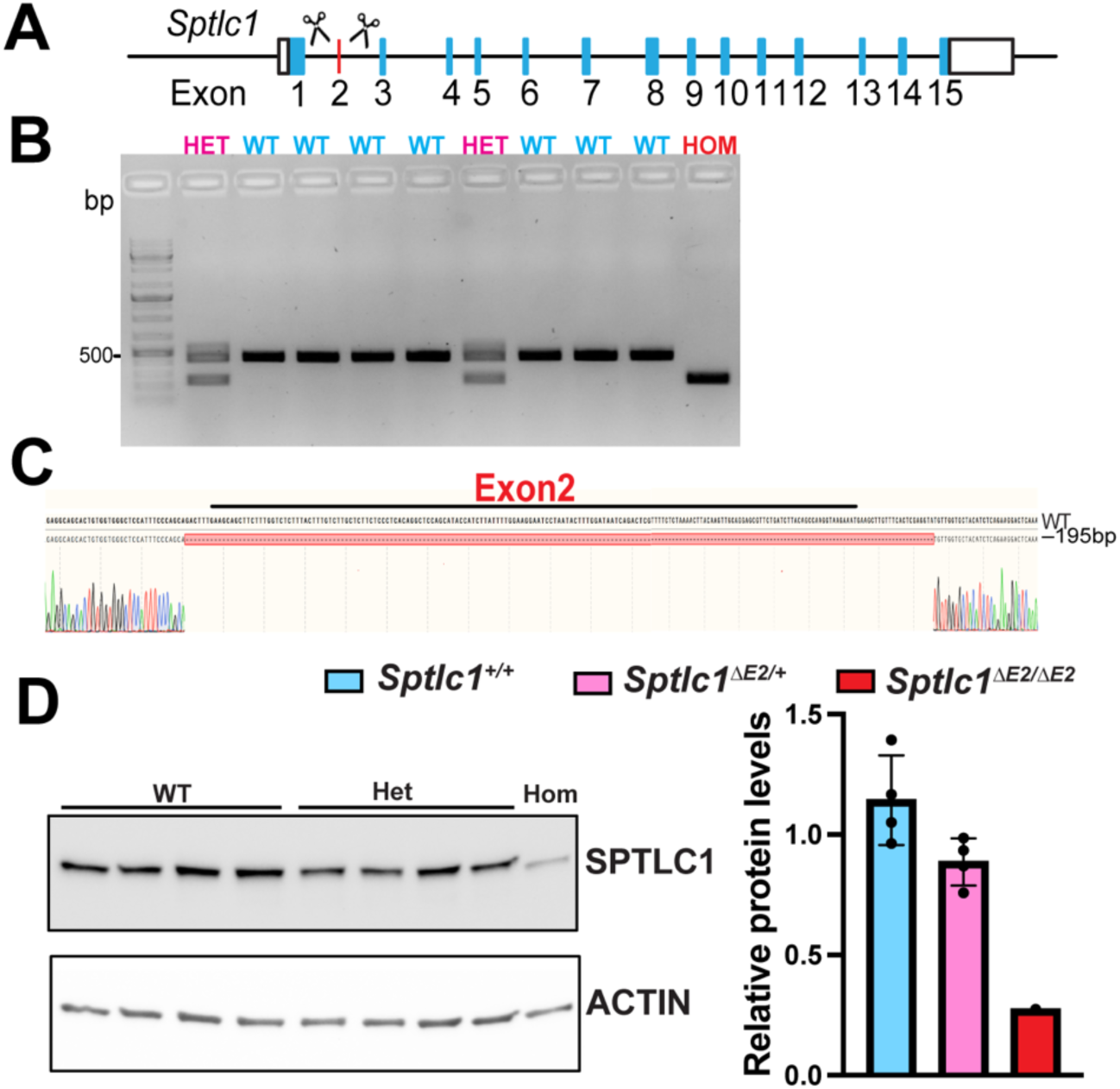
Generation of *Sptlc1* ΔE2 knock-in mice using CRISPR-Cas9 technology. **(A)** Gene diagram of the mouse *Sptlc1* locus showing the location of the exon 2 deletion. **(B)** Agarose gel electrophoresis with genotyping primers demonstrates precise deletion of exon 2 using the CRISPR-Cas9 knock-in approach. The heterozygous (HET) mouse shows two bands: the upper wildtype (WT) and a lower ΔE2 band, while the homozygous (HOM) mouse exhibits only the ΔE2 band. (**C**) Sanger sequencing of the purified PCR product from the HET animal confirms the *Sptlc1* wild type (WT) and exon 2 deletion (−195bp) sequences. (**D**) Western blot analysis shows no significant difference in total SPTLC1 protein levels between *Sptlc1*^+/+^ and *Sptlc1*^ΔE2/+^ spinal cords. Lysates from n = 4 mice per group were analyzed, and SPTLC1 levels were normalized to ACTIN. *Sptlc1*^ΔE2/ΔE2^ were lethal at an early stage and exhibited very low levels of total SPTLC1 protein.

### No increase in canonical sphingolipid levels in *Sptlc1^ΔE2/+^* mice

SPTLC1 is a core component of the SPT complex, which catalyzes the first step in sphingolipid biosynthesis by conjugating L-Serine with palmitoyl-CoA to generate sphingosine, the precursor to ceramide and complex sphingolipids. Given that ALS-associated *SPTLC1* variants have been proposed to disrupt sphingolipid homeostasis, we sought to determine the effect of exon 2 deletion on total sphingolipid amounts *in vivo*. Using mass spectrometry, we quantified canonical sphingolipids in both CNS and peripheral tissues of *Sptlc1^ΔE2/+^* mice and littermate controls at 4 months of age. Surprisingly, heterozygous *Sptlc1^ΔE2/+^* mice did not exhibit any significant alterations in sphingolipid levels across multiple tissues (**Fig. 2**). These findings suggest that the *Sptlc1* ΔE2 mutation does not impair bulk sphingolipid metabolism under basal conditions.

**Figure 2:**
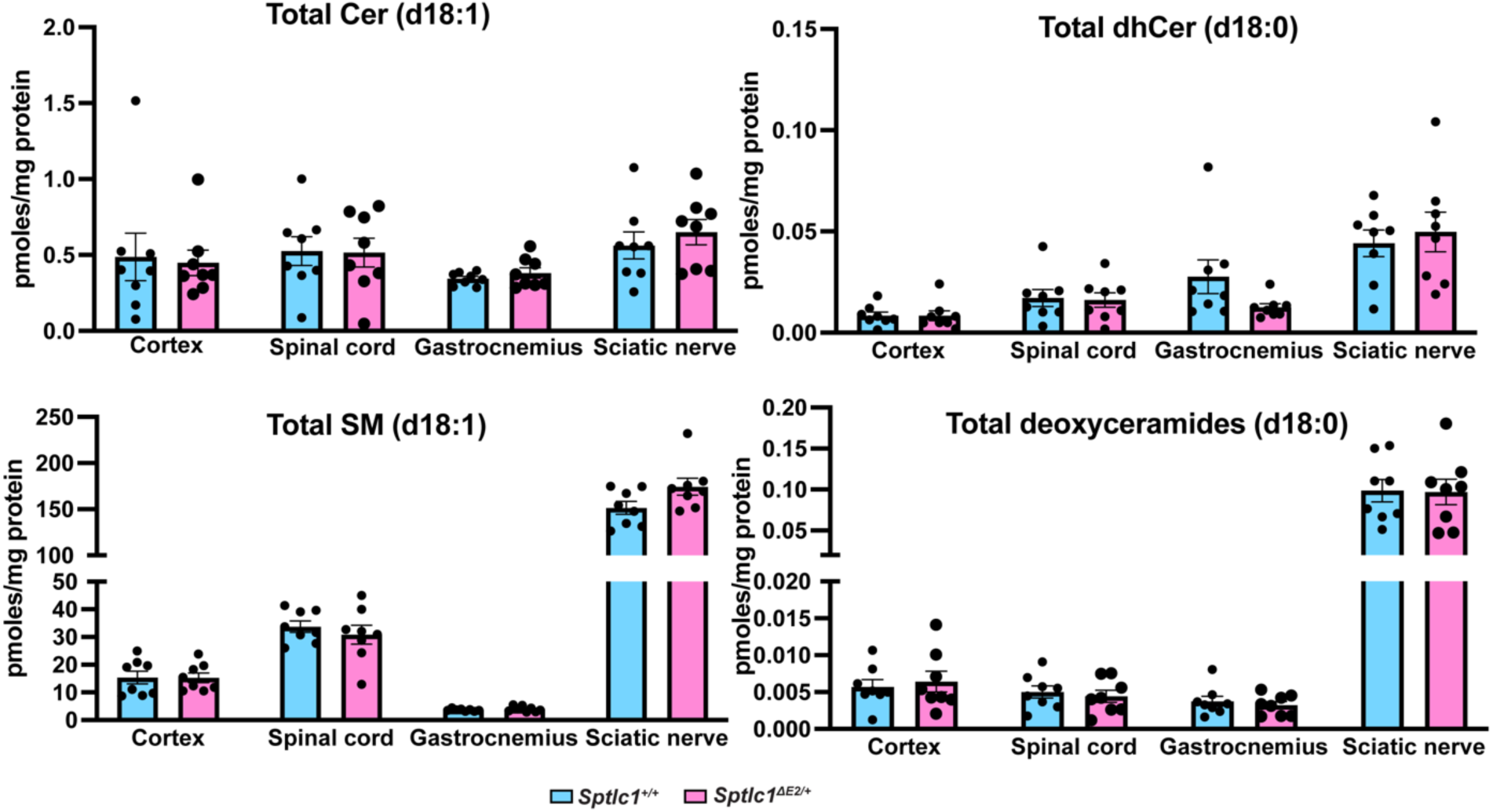
No alterations in lipidomics profile in *Sptlc1^ΔE2/+^* mice tissues. Similar profiles of canonical and non-canonical sphingolipids were observed in different mice tissues (cortex, spinal cord, gastrocnemius and sciatic nerve) in *Sptlc1*^ΔE2/+^ compared to controls or *Sptlc1*^+/+^. Data points are individual mice. Values are mean ± SEM (p<0.05, One-Way ANOVA, n=8 per group).

### *Sptlc1^ΔE2/+^* mice do not develop impaired motor functions

To test the effects of the ΔE2 mutation on motor function, we evaluated mice using various motor and behavioral tests at different time points. Male and female mice were tested on the rotarod assay beginning at 3 months of age, with no significant differences observed between *Sptlc1^+/+^* and mice up to 15 months of age (**Fig. 3A**). To further assess motor performance, we conducted grip strength assays through 15 months of age. No significant differences were detected between groups at any time point, based on the average performance across three trials (**Fig. 3B**). Additionally, muscle electrophysiology tests revealed no differences in compound muscle action potential (CMAP) amplitude in the hind paw muscles at 17 months of age (**Fig. 3C**). Overall, behavioral analyses did not identify any motor impairments in *Sptlc1^ΔE2/+^*mice. In addition, the heterozygotes *Sptlc1^ΔE2/+^* mice displayed no differences in survival and body weight compared to controls (data not shown), suggesting that the heterozygous ΔE2 mutations does not lead to major motor or neuromuscular dysfunction.

**Figure 3:**
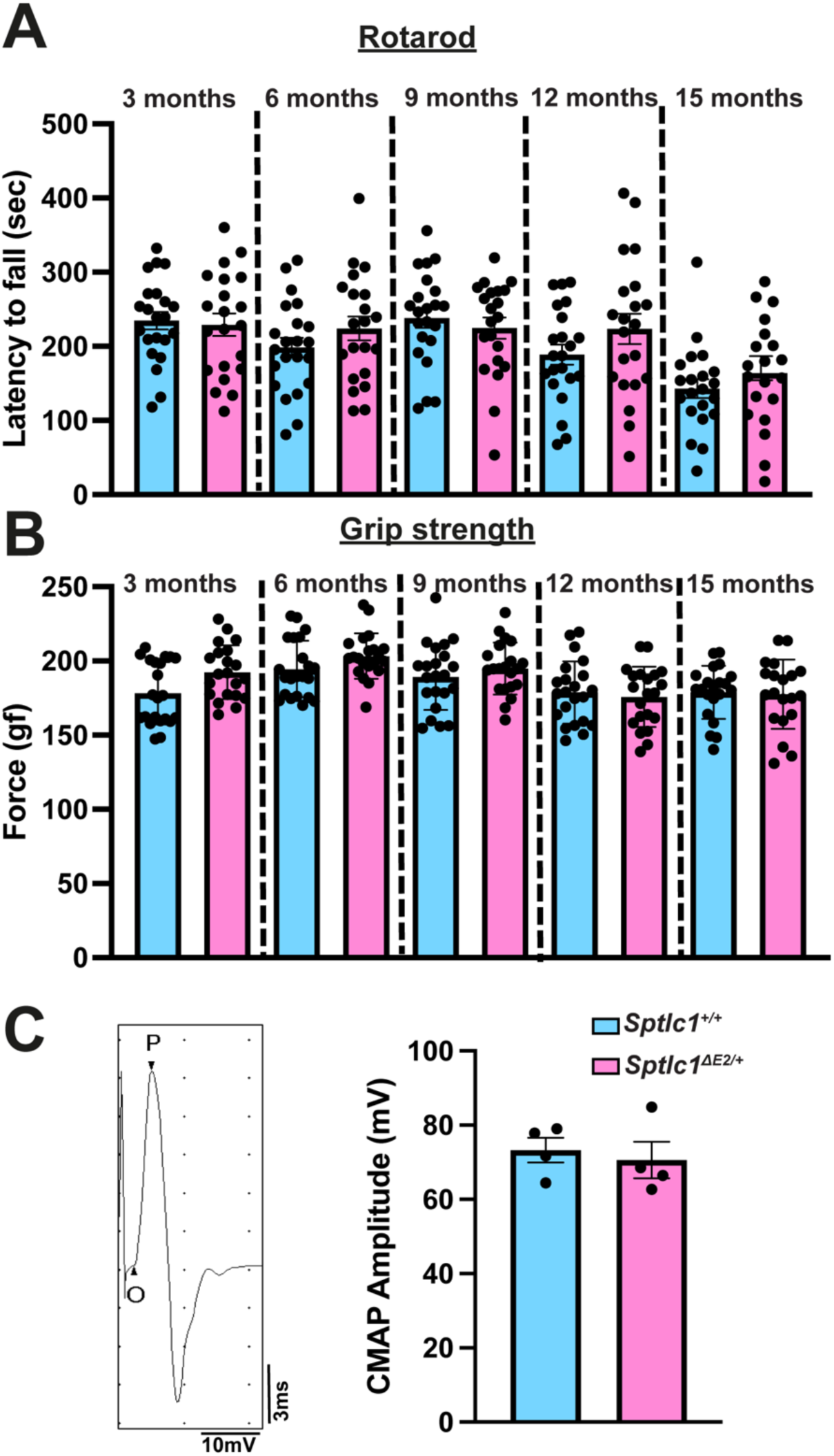
*Sptlc1^ΔE2/+^* mice do not display motor impairments. **(A)** Longitudinal rotarod analysis revealed that *Sptlc1*^ΔE2/+^ mice, both males and females, do not decline with age or significant changes compared to WT or *Sptlc1*^+/+^ littermates. **(B)** Muscle strength was assessed using a grip strength test and *Sptlc1*^ΔE2/+^ mice performed similarly compared to *Sptlc1*^+/+^ (p<0.05, One-Way ANOVA, n=18–20 per group. **(C)** Compound muscle action potential amplitudes (CMAP) in the hind paw muscles, elicited from nerve stimulation at the ankle, showed no differences in 18-month-old animals (p<0.05, student t test, n=4 mice per group).

### *Sptlc1^ΔE2/+^* mice do not display obvious neuroanatomical abnormalities

Magnetic resonance imaging (MRI) is a well-established tool in both clinical practice and neurological research. This noninvasive *in vivo* imaging modality offers a significant advantage by enabling translational MRI studies in rodent models of CNS diseases. MRI is known for its high reproducibility and minimal standard deviation, making it a reliable method for detecting structural brain changes [19]. To assess neuroanatomical alterations associated with the mutant SPTLC1 ΔE2, we performed *in vivo* MRI on 17-month-old heterozygous and wild-type littermates. Structural MRI demonstrated that most brain regions in mutant mice appeared comparable to controls (**Fig. 4A-B**). The analysis further revealed no statistically significant differences in total brain volume between heterozygous mutants and controls (**Fig. 4C**). These findings suggest that, at least at the level of gross anatomical structure, heterozygous *Sptlc1^ΔE2/+^* mice do not exhibit any overt neuroanatomical abnormalities.

**Figure 4.**
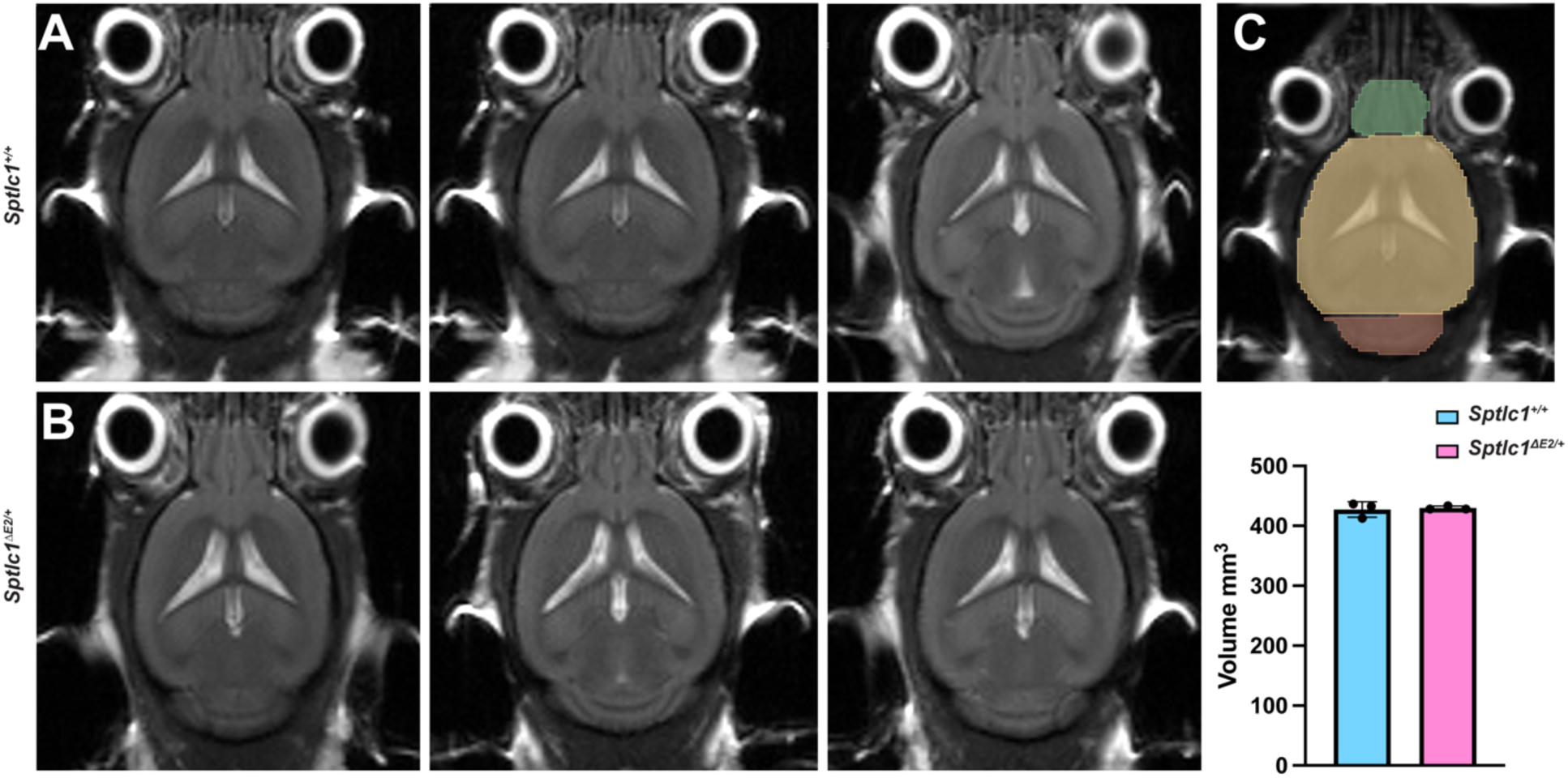
*Sptlc1^ΔE2/+^* mice brain MRI scans show no neuroanatomical abnormalities. *In vivo* MRI mouse brain images of *Sptlc1*^+/+^ (**A**) and *Sptlc1*^ΔE2/+^ one axial image (**B**). The mouse brain MRI image was extracted from a 2–dimensional dataset with the following parameters: FOV, 20 × 20 mm^2^; matrix, 116 × 116. (**C**) Brain structural segmentation, shown in color, was superimposed. The area was measured using 3D slicer (p<0.05, student t test, n=3 mice per group).

### *Sptlc1^ΔE2/+^* mice do not display neuronal loss or gliosis in the central nervous system

We next performed a comprehensive neuropathological analysis to determine the effects of the ΔE2 mutation. Immunofluorescence staining for NeuN, IBA1, and GFAP was performed on brain and lumbar spinal cord sections from 18-month-old mice. No obvious loss of neurons was observed in either brain (**Fig. S4**) or spinal cord (**Fig. 5A,B**). Additionally, there were no significant changes in astrocyte or microglial activation, as GFAP and IBA1 levels remain unaltered. Taken together, these findings indicate that heterozygous *Sptlc1^ΔE2/+^* mice do not display any noticeable cellular or molecular indicators of neurodegeneration or glial dysfunction.

**Figure 5.**
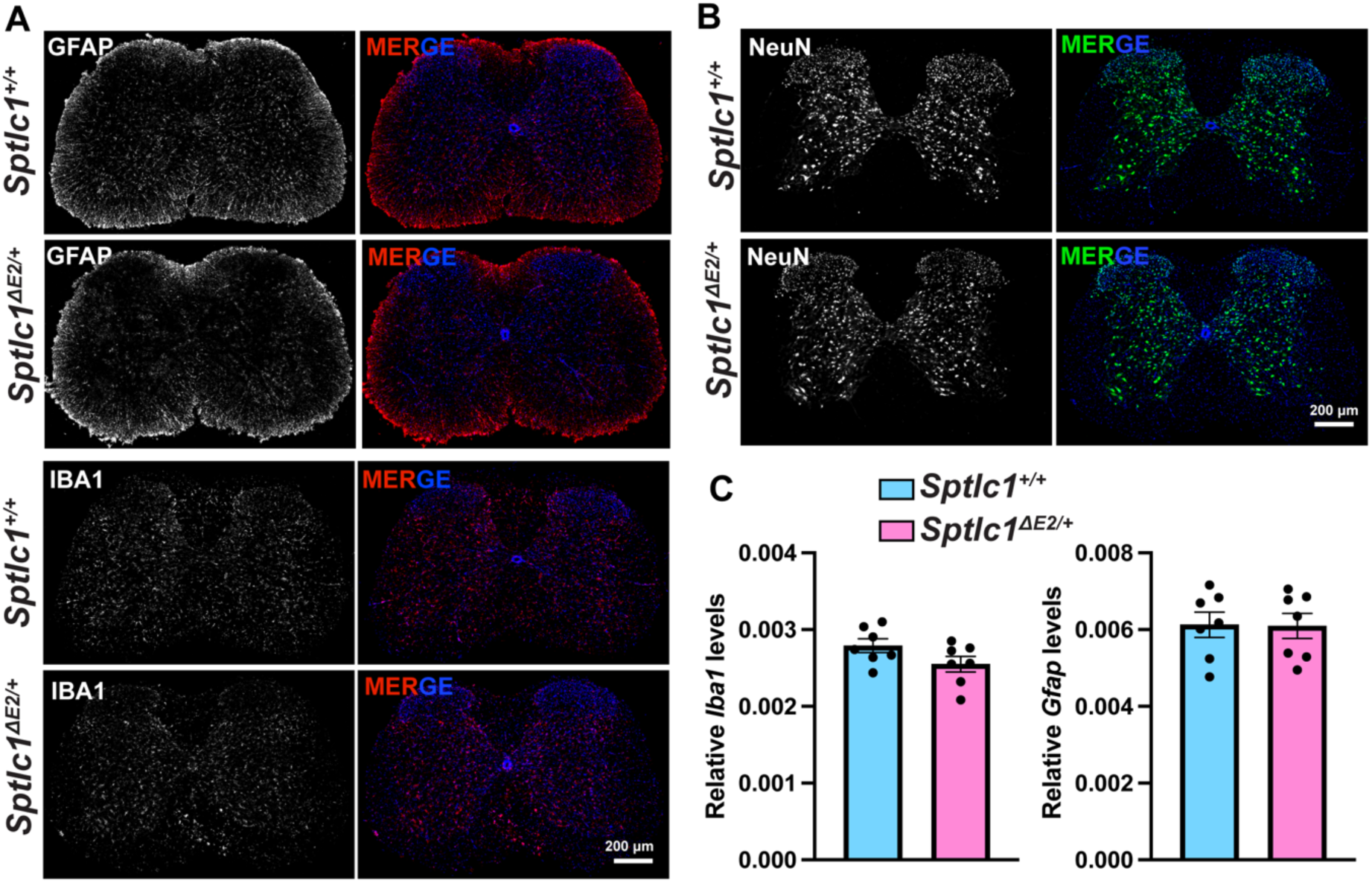
No evidence of reactive gliosis and neuronal loss in the *Sptlc1^ΔE2/+^* mice. GFAP and IBA1 immunofluorescence staining in 18 months old brains sagittal section (**A**) and lumbar spinal cord sections (**B**). The GFAP and IBA1 signals in *Sptlc1*^ΔE2/+^ were compared to wildtype littermates. Higher magnification images shown. NeuN was used as a neuronal marker. DAPI was used to stain nuclei. N=3 mice per group. (**C**) RT-PCR confirms no significant difference in *Gfap* or *Iba1* levels in cortex and spinal cord regions of *Sptlc1*^ΔE2/+^ compared to controls or *Sptlc1*^+/+^ 18 months old animals (p<0.05, student t test, n=7 mice per group).

### *Sptlc1^ΔE2/+^* mice exhibit no detectable changes in SPT complex

Recent structural studies have provided key insights into the organization and regulation the SPT complex. SPTLC1 and SPTLC2 form the catalytic core of the enzyme, while ORMDL proteins act as negative regulators by modulating SPT activity in response to sphingolipid levels. Given that SPTLC1-linked jALS variants cluster near the transmembrane domain, potentially affecting protein-protein interactions within the complex, we investigated whether the *Sptlc1* ΔE2 mutation alters the expression of key SPT components *in vivo*. Due to the lack of commercial antibodies for mouse SPTLC2 and ORMDL3, we quantified transcript levels using qPCR. Expression analysis revealed no significant changes in *Sptlc1, Sptlc2, Ormdl3* mRNA levels in *Sptlc1^ΔE2/+^*mice compared to littermate controls (**Fig. 6**). These findings suggest that the *Sptlc1* ΔE2 mutation does not disrupt the expression of SPT complex components.

**Figure 6.**
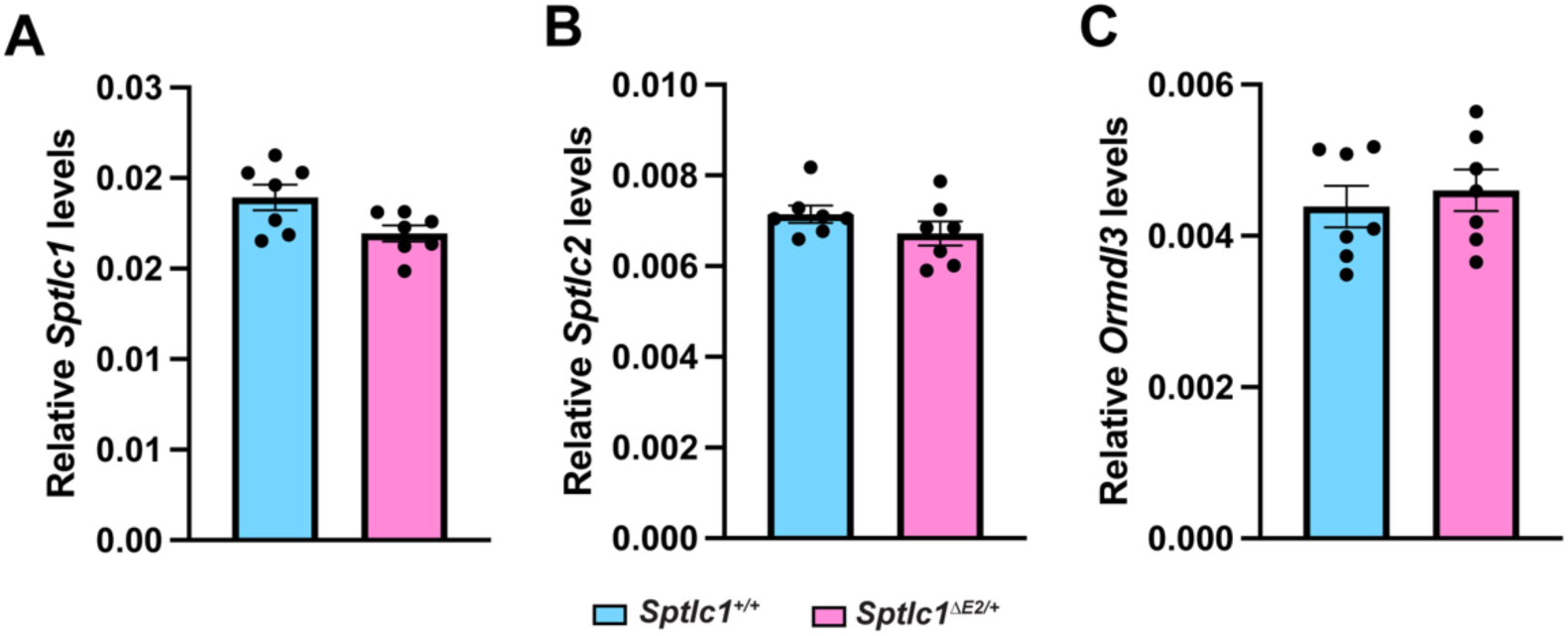
*Sptlc1^ΔE2/+^* did not induce any changes in the SPT-ORMDL complex. **(A)** Total *Sptlc1* mRNA levels were measured using exon 3-4 junction probes. (**B-C**) *Sptlc2* and *Ormdl3* mRNA levels were measured using exon 4-5 junction probes. Data are presented as mean ± SEM, p<0.05, student t test, n=7 mice per group.

## Discussion

In this study, we generated and characterized a novel *Sptlc1* exon 2 deletion knock-in mouse model to investigate the pathogenesis of SPTLC1-associated jALS. While prior studies have suggested that SPTLC1-jALS variants lead to dysregulated sphingolipid metabolism, our findings indicate that heterozygous (*Sptlc1^ΔE2/+^*) mice do not exhibit significant disruptions in sphingolipid homeostasis, motor function, neuroanatomy, or neuropathology.

To date, missense mutations in *SPTLC1* have been linked several human diseases including ALS, HSAN1 and Flegel disease [5, 10, 20]. The SPTLC1-HSAN1 linked mutations typically result in a loss of substrate specificity, causing the SPT enzyme to incorporate alanine or glycine instead of serine to produce deoxysphingolipids. Deoxysphingolipids, which lack a critical hydroxyl moiety, cannot be efficiently degraded by the cell machinery and cause toxicity. In contrast, ALS-associated *SPTLC1*, including Y23F, L38R, L39del and F40S41del, are clustered in exon 2 and one of them, c.58G/T, leads to exon 2 skipping [5, 21]. Exon 2 of SPTLC1 forms a transmembrane domain that interacts with ORMDL proteins, negative regulator of SPT activity. By overexpressing the ALS-associated *SPTLC1* mutations in HEK293 cells with endogenous *SPTLC1* deleted, Lone et al. showed that these variants impaired its interaction with ORMDLs, resulting in increased sphingolipid synthesis and a distinct lipid signature [16]. Canonical sphingolipids were also shown to increase in human iPSC derived motor neurons using a compound heterozygous (*SPTLC1^ΔF40_S41/ΔE2^*) line [5]. However, the ALS patient reported so far doesn’t have any compound heterozygous mutation. Therefore, an increase in sphingolipids in this model might be driven by a different mechanism.

Despite this *in vitro* evidence, we found no significant alterations in canonical sphingolipid levels in CNS or peripheral tissues of *Sptlc1*^ΔE2/+^ mice. One possible explanation for this discrepancy is that there are compensatory mechanisms in mice for the ΔE2 deletion. While SPTLC1 protein with ΔE2 deletion are less stable as seen in homozygote mice, the total level of SPTLC1 is not changed in the *Sptlc1*^ΔE2/+^ mice. Alternatively, the impact of ALS-associated *SPTLC1* mutations on lipid metabolism may be more context-dependent than previously thought. Studies measuring sphingolipid levels in patient-derived biofluids have produced conflicting results. Mohassel *et al,* showed that c.56G/T patient plasma had increased formation of canonical sphingolipids whereas another study from Johnson *et al*, showed no change in sphingolipids levels in two different patients with same mutation [3, 5]. Given that lipid metabolism is influenced by multiple factors, including age, diet, and systemic metabolic regulation, our findings highlight the need for further investigation into the specific conditions under which *SPTLC1* ΔE2 variant might lead to dysregulated sphingolipid synthesis.

A key observation in our study is the embryonic lethality associated with the homozygous *Sptlc1* ΔE2 allele, albeit with incomplete penetrance. Due to early lethality of our homozygous ΔE2 mice, it hinders our efforts to generate a viable ΔE2 homozygous mice cohort to perform the experiments together with heterozygous or control mice. The *Sptlc1^ΔE2^* knockin allele is informative. While *Sptlc1* knockout mice were reported embryonic lethal, therefore, it is surprising that the ΔExon2 allele is not viable when homozygous. It is possible that very low levels of the total SPTLC1 protein in these mice would results in aberrant sphingolipids homeostasis during development and would reflect a sharp threshold when transitioning to a lethal phenotype due to loss-of-function. Alternatively, unstable stoichiometry of SPT complex subunits (SPTLC1, SPTLC2, ORMDL3) may be irreparably perturbed in homozygotes. In a human patient with exon 2 skipping variant, proteomics studies revealed a 50% reduction in total SPTLC1 levels in lymphoblast cells [21]. However, there was no lipidomic study done in patient cells [21].

The first mouse models of *Sptlc1* to understand the role of *SPTLC1* in human disease context used overexpression approach. Overexpression of SPTLC1-linked HSAN1 C133W mutation transgenic mice (using chicken beta-actin promoter with CMV enhancer elements and *Cricetulus griseus* SPTLC1 cDNA) displayed age-dependent weight loss and mild sensory and motor impairments [13]. Similarly, overexpression of fusion SPT (fSPT) construct consisting of three individual wild type SPT subunits (SPTLC2, SPTSSA, and SPTLC1) results in neurological phenotype in mice [22]. Together, these studies demonstrate how increased enzyme activity levels can lead to severe phenotypes in rodents. Although these models were informative but present a caveat that perhaps the pathology observed in these models is due to transgene overexpression, and not reflective of the human disease state that exhibits endogenous gene expression. Recently, knock-in *Sptlc1*^C133W^ mouse model have been generated to address this for HSN1 disease by Burgess laboratory [12]. The heterozygous mice were normal and displayed biochemical phenotype of HSAN1 disease and mice were lethal as homozygotes. It is not clear why homozygotes *Sptlc1*^C133W^ were lethal. Nevertheless, this data suggests that sphingolipid synthesis is tightly controlled and are critical for mouse development. Collectively, we demonstrate that *Sptlc1^ΔE2/+^* knock-in mice do not recapitulate ALS features such as motor function impairment. It is also possible that additional genetic factors may influence disease progression. Nonetheless, this model has been informative and will be useful for future mechanistic studies.

## Acknowledgements

The authors would like to thank the members of the Jiang and Hornemann labs for their support. We also extend our gratitude to Emory ALS Center for their helpful discussions. This study was supported in part by the Mouse Transgenic and Gene Targeting Core (TMF), which is subsidized by the Emory University School of Medicine and is part of the Emory Integrated Core Facilities. Additional support was provided by the Emory University Center for Systems Imaging Core Facility (RRID:SCR_023522) and by the National Center for Advancing Translational Sciences of the National Institutes of Health (NIH) under award number UL1TR000454. The content is solely the responsibility of the authors and does not necessarily reflect the official views of the NIH. This work was funded by the Safenowitz Postdoctoral Fellowship from the ALSA (22-PDF-605) and Career Development Award (78095) from the LLLF to DCP. Additionally, JP, FM, JJ are partially supported by the NIH R01 grant (R01AG068247) awarded to JJ.

