## Supplementary figs for "Lack of motor defects and ALS-like neuropathology in heterozygous *Sptlc1* Exon 2 deletion mice"

Supplementary figures

```
CLUSTAL O(1.2.4) multiple sequence alignment

                                      TM (Exon 2)

hSPTLC1      MATATEQWVLVEMVQALYEAPAYHLILEGILILWIIRLLFSKTYKLQERSDLTVKEKEEL      60
mSPTLC1      MATVAEQWVLVEMVQALYEAPAYHLILEGILILWIIRLVFSKTYKLQERSDLTAKEKEEL      60
             ***.:*********************************:**************.******

hSPTLC1      IEEWQPEPLVPPVPKDHPALNYNIVSGPPSHKTVVNGKECINFASFNFLGLLDNPRVKAA      120
mSPTLC1      IEEWQPEPLVPPVSKNHPALNYNIVSGPPTHNIVVNGKECVNFASFNFLGLLANPRVKAT      120
             ************* *:*************:*: *******:*********** ******:

hSPTLC1      ALASLKKYGVGTCGPRGFYGTFDVHLDLEDRLAKFMKTEEAIIYSYGFATIASAIPAYSK      180
mSPTLC1      AFSSLKKYGVGTCGPRGFYGTFDVHLDLEERLAKFMKTEEAIIYSYGFSTIASAIPAYSK      180
             *::**************************:******************:***********

hSPTLC1      RGDIVFVDRAACFAIQKGLQASRSDIKLFKHNDMADLERLLKEQEIEDQKNPRKARVTRR      240
mSPTLC1      RGDIIFVDSAACFAIQKGLQASRSDIKLFKHNDVADLERLLKEQEIEDQKNPRKARVTRR      240
             ****:*** ************************:**************************

hSPTLC1      FIVVEGLYMNTGTICPLPELVKLKYKYKARIFLEESLSFGVLGEHGRGVTEHYGINIDDI      300
mSPTLC1      FIVVEGLYMNTGTICPLPELVKLKYKYKARIFLEESLSFGVLGEHGRGVTEHYGISIDDI      300
             *******************************************************.****

hSPTLC1      DLISANMENALASIGGFCCGRSFVIDHQRLSGQGYCFSASLPPLLAAAAIEALNIMEENP      360
mSPTLC1      DLISANMENALASVGGFCCGRSFVVDHQRLSGQGYCFSASLPPLLAAAAIEALNIMEENP      360
             *************:**********:***********************************

hSPTLC1      GIFAVLKEKCGQIHKALQGISGLKVVGESLSPAFHLQLEESTGSREQDVRLLQEIVDQCM      420
mSPTLC1      DIFAVLKKKCQNIHKSLQGVSGLKVVGESLSPALHLQLEESTGSREKDVKLLQAIVDQCM      420
             .******:** :***:***:*************:************:**:*** ******

hSPTLC1      NRSIALTQARYLEKEEKCLPPPSIRVVVTVEQTEEELERAASTIKEVAQAVLL      473
mSPTLC1      DKGIALTQARYLDKEEKCLPPPSIRVVVTVEQTEEELQRAASTIREAAQAVLL      473
             ::.*********:************************:******:*.******
```

**Figure S1. Overall organization of mouse and human SPTLC1.**
SPTLC1 comprised of transmembrane (TM) domain encoded by exon 2. Protein sequence alignment of human (NP_006406.1), mouse (NP_006406.1), showing position of highly conserved TM domain (red box). The c.58G/T mutation that results in skipping of exon 2 result in a short sequence from residue 20 onward.

| F0 # | Sex | Indel (bp) | gRNA + Donor |
|---|---|---|---|
| 34 | M | -37 | 2 guides |
| 35 | M | -126 | 2 guides |
| 36 | M | 0 | 2 guides |
| 37 | M | 0 | 2 guides |
| 38 | M | -20 | 2 guides |
| 39 | M | 0 | 2 guides |
| 40 | F | 0 | 2 guides |
| 41 | F | -195 | 2 guides |
| 42 | F | 0 | 2 guides |
| 43 | F | -6 | 2 guides |
| 44 | F |  | 2 guides |
| 45 | F | -8 | 2 guides |
| 46 | M | 0 | 2 guides |
| 47 | M | -196 | 2 guides |
| 48 | M | 0 | 3 guides |
| 49 | M | 0 | 3 guides |
| 50 | M | 0 | 3 guides |
| 51 | M | 0 | 3 guides |
| 52 | F | 0 | 3 guides |
| 53 | F | -20 | 3 guides |
| 54 | F | 0 | 3 guides |
| 55 | F | 0 | 3 guides |
| 56 | M | -195 | 2 guides |
| 57 | M | -195 | 2 guides |
| 58 | M | -44 | 2 guides |
| 59 | M | 0 | 2 guides |
| 60 | F | 0 | 2 guides |
| 61 | F | -8 | 2 guides |
| 62 | F | 0 | 2 guides |
| 63 | F | 0 | 2 guides |
| 64 | F | 0 | 2 guides |
| 65 | M | 0 | 2 or 3 guides |
| 66 | M | 0 | 2 or 3 guides |
| 67 | M | 0 | 2 or 3 guides |
| 68 | M | 0 | 2 or 3 guides |
| 69 | M | 0 | 2 or 3 guides |
| 70 | M | 0 | 2 or 3 guides |
| 71 | M | 0 | 2 or 3 guides |
| 72 | M | 0 | 2 or 3 guides |
| 73 | M | 0 | 2 or 3 guides |
| 74 | F | 0 | 2 or 3 guides |
| 75 | F | 0 | 2 or 3 guides |
| 76 | F | 0 | 2 or 3 guides |
| 77 | M | 0 | 2 guides |
| 78 | M | 0 | 2 guides |
| 79 | M | 0 | 2 guides |
| 80 | M | -8 | 2 guides |
| 81 | M | -121 | 2 guides |
| 82 | M | 0 | 2 guides |
| 83 | M | -20 | 2 guides |
| 84 | F | 0 | 2 guides |
| 85 | F | 0 | 2 guides |
| 86 | F | 0 | 2 guides |
| 87 | F | 0 | 2 guides |
| 88 | F | -45 | 2 guides |
| 89 | F | 0 | 2 guides |
| 90 | M | 0 | 3 guides |
| 91 | M | 0 | 3 guides |
| 92 | M | 0 | 3 guides |
| 93 | F | 0 | 3 guides |
| 94 | F | 0 | 3 guides |
| 95 | F | 0 | 3 guides |

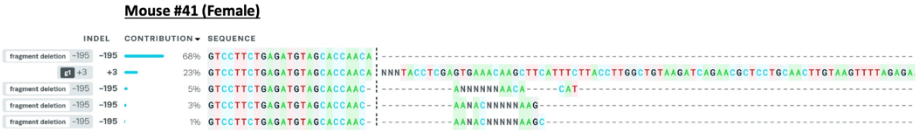


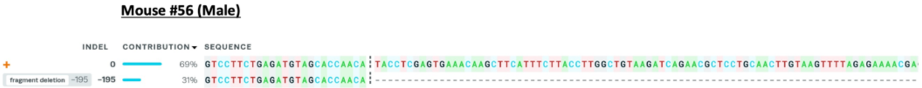


**Figure S2. Analysis of sgRNAs that target the introns spaning *Sptlc1* exon 2 locus.**
Sequences of the 64 mice injected with Cas9, sgRNAs and donor oligo shows cleavage of the *Sptlc1* locus at the intron-exon-intron junction of exon 2. Base modification status are listed. Two positive mosaic founders (F0) lines #41, #56 (in red) were selected as they displayed precise deletion of exon2 with highest % indel contribution (-195bp) as confimed by Synthego ICE in-silico analysis.

**A**

ATGGCGACAGTGGCGGAGCAGTGGGTG**<u>CTGGTGGAGATGGTGCAGG</u>**CGCTGTACGAGGCTCCAG
CATACCATCTTATTTTGGAAGGAATCCTAATACTTTGGATAATCAGACTCGTTTTCTCTAAAACTTACAA
GTTGCAGGAGCGTTCTGATCTTACAGCCAAGGAAAAGGAAGAACTGATTGAAGAGTGGCAGCCAGA
GCCCCTCGTCCCTCCAGTCTCCAAGAACCACCCTGCTCTCAACTACAACATCGTGTCCGGCCCTCC
AACCCACAACATCGTGGTGAATGGAAAAGAGTGTGTCAACTTTGCCTCCTTTAACTTCCTTGGGCTG
CTGGCCAACCCTCGAGTTAAGGCCACAGCTTTTTCATCTTTAAAGAAGTACGGAGTGGGTACCTGTG
GTCCT**<u>CGAGGGTTCTATGGCACATTTG</u>**

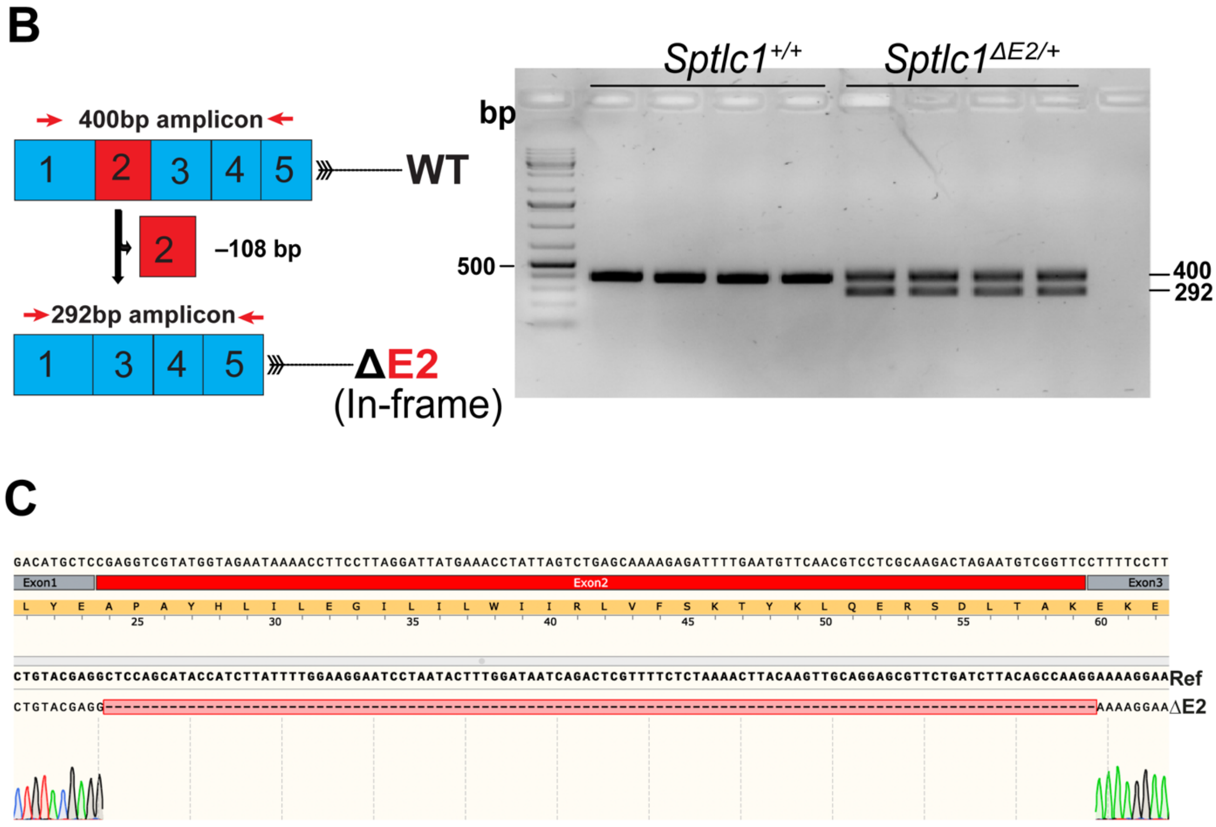


**Figure S3. Characterization of the *Sptlc1*$^{ΔE2/+}$ mouse F1 line.**
(**A**) cDNA sequence of targeted locus of *Sptlc1* exon 2 and surrounding exon 2 regions. Primer sequences in exon 1 (forward) and exon 5 (reverse) bold and underlined, exon 2 indicated in red. (**B**) In-frame deletion of exon 2. Agarose gel electrophoresis of RT-PCR products to validate deletion of exon 2 using mRNA from mice spinal cord tissues. RT-PCR primers were in exons 1 and 5 (red arrow), and the amplicon size 400 bp for WT mice and 292 bp for ΔE2 F1 mice (n=4, 4 m old). (**C**) Sanger sequencing confirmed the precise deletion of only exon 2.

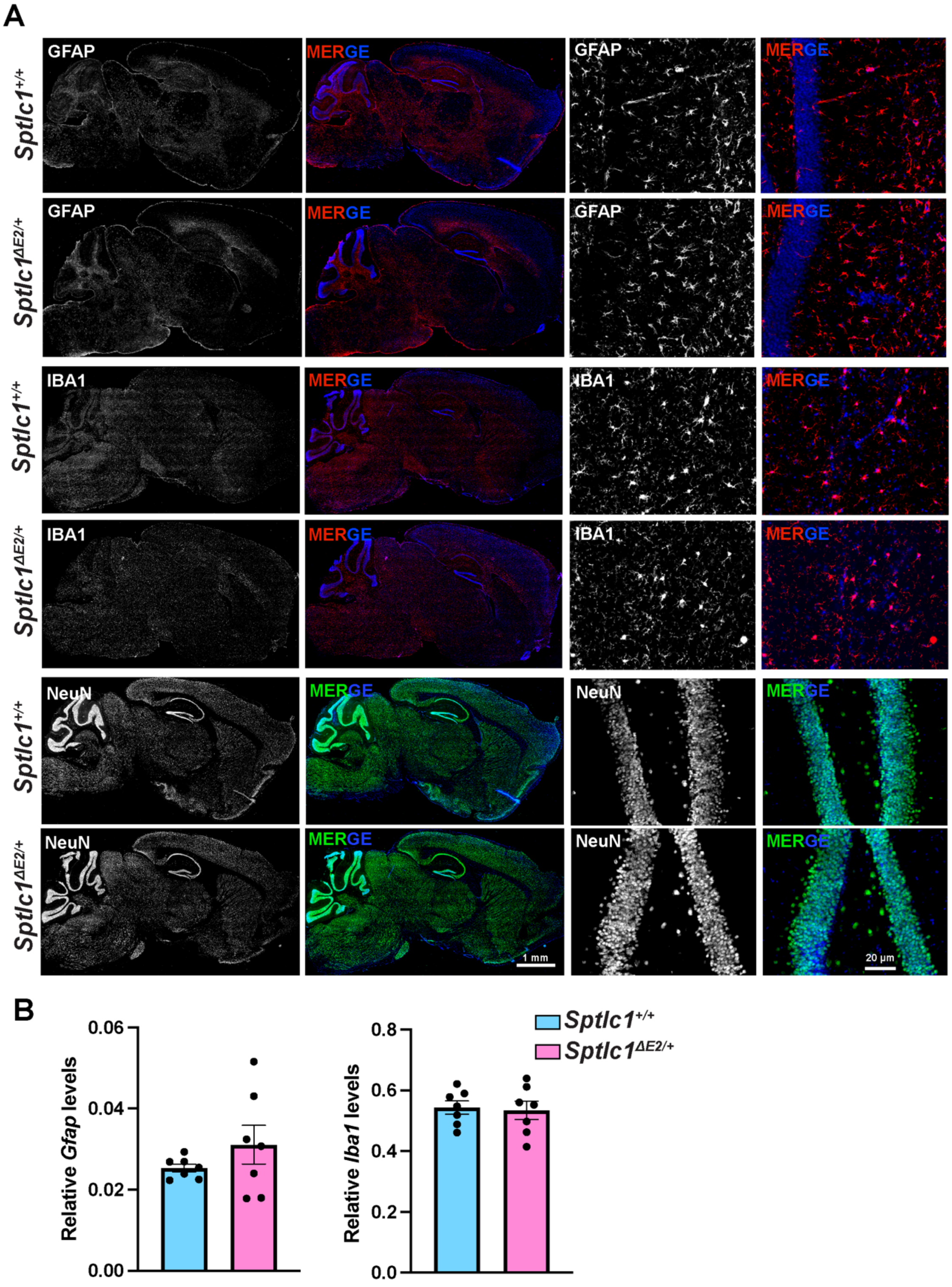


**Figure S4. No evidence of reactive gliosis and neuronal loss in the *Sptlc1*$^{\Delta E2/+}$ mice brain.**
(**A**) GFAP and IBA1 immunofluorescence staining in 18 months old brains sagittal section. The GFAP and IBA1 signals in *Sptlc1*$^{\Delta E2/+}$ were compared to wildtype littermates. Higher magnification images shown. NeuN was used as a neuronal marker. DAPI was used to stain nuclei. N=3 mice per group. (**B**) RT-PCR confirms no significant difference in Gfap or Iba1 levels in cortex region of *Sptlc1*$^{\Delta E2/+}$ compared to controls or *Sptlc1*$^{+/+}$ 18 months old animals ($p<0.05$, student t test, n=7 mice per group).
